# Plasmid-borne *mcr-1* and Replicative Transposition of Episomal and Chromosomal blaNDM-1, blaOXA-69, and blaOXA-23 Carbapenemases in a Clinical *Acinetobacter baumannii* Isolate

**DOI:** 10.1101/2024.08.27.609937

**Authors:** Masego Mmatli, Nontombi Marylucy Mbelle, John Osei Sekyere

**Affiliations:** Department of Medical Microbiology, School of Medicine, University of Pretoria, South Africa; Department of Medical Microbiology, Tshwane Academic Division, National Health Laboratory Service, Pretoria, South Africa; Department of Dermatology, School of Medicine, University of Pretoria, South Africa

**Keywords:** colistin resistance, carbapenem, carbapenemase, last-resort antibiotics, non-fermenters, multi-drug resistance, RNAseq

## Abstract

**Background:** A multidrug-resistant clinical A. baumannii isolate with resistance to most antibiotics was isolated from a patient at an intensive care unit. The genetic environment, transcriptome, mobile, and resistome were characterized.

**Method:** The MicroScan system, disc diffusion, and broth microdilution were used to determine the resistance profile of the isolate. A multiplex PCR assay was also used to screen for carbapenemases and mcr-1 to -5 resistance genes. Efflux-pump inhibitors were used to evaluate efflux activity. The resistome, mobilome, epigenome, and transcriptome were characterized.

**Results & conclusion:** There was phenotypic resistance to 22 of the 25 antibiotics tested, intermediate resistance to levofloxacin and nalidixic acid, and susceptibility to tigecycline, which corresponded to the 27 resistance genes found in the genome, most of which occurred in multiple copies through replicative transposition. A plasmid-borne (pR-B2.MM_C3) *mcr-*1 and chromosomal *bla_PER-7_, bla_OXA-69_, bla_OXA-23_* (three copies), *bla_ADC-25_*, *bla_TEM-1B_*, and *bla*_NDM-1_ were found within composite transposons, ISs, and/or class 1 and 2 integrons on genomic islands. Types I and II methylases and restriction endonucleases were in close synteny to these resistance genes within the genomic islands; chromosomal genomic islands aligned with known plasmids. There was a closer evolutionary relationship between the strain and global strains but not local or regional strains; the resistomes also differed. Significantly expressed/repressed genes (6.2%) included resistance genes, hypothetical proteins, mobile elements, methyltransferases, transcription factors, membrane and efflux proteins.

The genomic evolution observed in this strain explains its adaptability and pandrug resistance and shows its genomic plasticity on exposure to antibiotics.

## 1. Introduction

*Acinetobacter baumannii* is an aerobic, coccobacillary rod, non-fermenting Gram-negative pathogen ^1,2^. It is an important, ubiquitous and opportunistic pathogen found in both moist and dry conditions and is well distributed within nature, the nosocomial environment and the human mucosal microbiome ^3^. *A. baumannii* causes both community and health-care associated infections (HAIs) ^4,5^ including urinary tract, bloodstream, skin, and tissue, as well as ventilator-associated infections ^6,7^. These infections are usually difficult to treat and are fatal ^8,9^ as *A. baumannii* can survive for prolonged periods in the hospital environment, facilitating its nosocomial spread ^10^. This is achieved through either direct contact with an infected patient (patient-to-patient), contact with the hands of health care personnel, or indirectly by touching contaminated environmental surfaces ^1,3,11^. Individuals at risk for *A. baumannii*-related HAIs are typically those who are immuno-deficient ^7,10^, or are undergoing invasive procedures such as the use of mechanical ventilators, central venous or urinary catheters, as seen in the invasive care unit (ICU) ^4,10^.

Carbapenem resistance in *A. baumannii* is mainly acquired through the production of oxacillinase-type carbapenemases, with *bla*_OXA-23_-like and *bla*_OXA-48_-like being the most prevalent β-lactamase (carbapenemase) ^12,13^. Hence, carbapenem-resistant *A. baumannii* (CRAB) ^13^ is mostly treated using colistin and tigecycline as the last option. While phosphoethanolamine (PEtN)-mediated CRAB has been reported in South Africa ^14,15^, mobile colistin resistance (mcr) genes have been reported in *A. baumannii* in China ^16^, Brazil ^17^, Italy ^18^, Pakistan ^7^, Turkey ^5^, Europe 23 and Iraq 24. Nevertheless, there has been no reports of mcr-producing A. baumannii in South Africa. Hence, this study presents the first report of an *mcr-*positive CRAB isolate (with multiple carbapenemases) from South Africa and to our knowledge, the first globally, using genomics, transcriptomics, epigenomics, and advanced bioinformatics to characterize its mechanisms of resistance and genome structure.

## 2. Methods

### Sample source and phenotypic resistance

A 53-year-old female patient at an intensive care unit (ICU) of the Steve Biko Academic Hospital, a tertiary and quaternary hospital in Pretoria, South Africa, presented with a difficult-to-treat infection in 2017. A fluid aspirate from the patient was cultured on blood agar media for 24 hours at 37℃ and subsequently identified in the laboratory; the isolate was labelled as R-B2.MM ^19^. The Microscan Walkaway identification/antibiotic susceptibility testing system (Beckman Coulter Diagnostics, United States) with Panel Combo 68 was used to identify the species and antibiotic resistance profiles of the isolate. Carbapenem and colistin resistance of the isolate were confirmed using the disc diffusion (10 µg discs of ertapenem, meropenem and imipenem) and broth microdilution (BMD) methods, respectively. The Clinical Laboratory Standards Institute (CLSI) ^20^ and the European Committee on Antimicrobial Susceptibility Testing (EUCAST)^21^ breakpoints were used respectively for the non-colistin antibiotics and colistin BMD. The BMD assay was performed using colistin sulphate powder, according to the CLSI standards.

Ertapenem sulphate salt and colistin sulphate salt (Glentham Life Sciences, United Kingdom), were used for the BMD assay ^22^. *Escherichia coli* ATCC 25922 and *Pseudomonas aeruginosa* ATCC 27853 were included as quality control strains. Both antibiotics were dissolved in sterile deionized water according to the manufacturers’ instructions. The antibiotic concentrations tested were: 128 µg/mL, 64 µg/mL, 32 µg/mL, 16 µg/mL, 8 µg/mL, 4 µg/mL, 2 µg/mL, 1 µg/mL, 0.5 µg/mL, and 0.25 µg/mL.

The BMD assay was performed in untreated 96-well polystyrene microtiter plates, with each well containing 100 µL of antibiotic dilution and Mueller-Hinton broth (MHB) or cation-adjusted MHB for ertapenem and colistin respectively. Subsequently, a 0.5 MacFarland suspension of bacterial culture was prepared, diluted to 1:20 with sterile saline, and 0.01 mL was inoculated into each well. The plates also included sensitive and negative control wells.

### Efflux-pump inhibitors

The role of efflux pumps in the resistance mechanisms of the isolate was investigated using the BMD method and the following efflux-pump inhibitors (EPIs): verapamil, phenylalanine-arginine β-naphthylamide (PAβN), carbonyl cyanide 3-chlorophenylhydrazone (CCCP), reserpine, and ethylenediaminetetraacetic acid (EDTA). The change in carbapenem and colistin resistance minimum inhibitory concentrations (MICSs) in the presence of the EPIs were calculated. The BMD and agar plates were incubated at 37 °C for 16-18 hours, and the minimum inhibitory concentration (MIC) was determined as the lowest antibiotic concentration without visible bacterial growth; the inhibition zones (for the disc diffusion tests) were used to determine carbapenem resistance using the CLSI breakpoints ^20,23^. The final concentrations of the antibiotic substrates in the broth were 1.5 µg/mL for CCCP, 4 µg/mL for VER, 25 µg/mL for PAβN, 20 µg/mL for RES, and 20 mM (pH 8.0) for EDTA. A ≥ 2-fold reduction in ertapenem and colistin MICs after EPI applications was indicative of significant efflux pump, metallo β-lactamase, and MCR activity and role in carbapenem or colistin resistance.

### Molecular characterization

Genomic DNA and RNA were respectively extracted from a 24-hour culture using Quick-DNA-fungal/bacterial MiniPrep™ kit (ZymoResearch) and Quick-RNA-fungal/bacterial MiniPrep™ kit (Zymo Research) according to the manufacturer’s protocols. Prior to RNA extraction, the bacterial suspension was grown in a broth containing 0.5 mg/mL of ertapenem and 2 mg/mL of colistin for at least 12 hours. The RNA was converted into cDNA using Qiagen’s cDNA synthesis kit.

Aliquots of the gDNA was used in a multiplex PCR screening test to identify the presence of *mcr* and carbapenemases using the primers in Table S1 (Dataset 1) and conditions already described in another study^19^. The gDNA was sequenced using PacBio SMRT sequencing at 100x coverage while the cDNA was sequenced using Illumina Miseq at a commercial sequencing facility. The PacBio reads (in fastQ format) were assembled using PacBio’s hierarchical genome-assembly process (HGAP) software to obtain a fastA file, which was annotated with NCBIs Prokaryotic Genome Annotation Pipeline (PGAP). The methylation files (motifs and base modifications) of the genome was also obtained from PacBio’s SMRTAnalysis MotifMaker software.

The species, MLST profile, resistome, mobilome, and epigenome, of the isolate were determined using NCBIs Average Nucleotide Identity (ANI) ^24^, Center for Genomic Epidemiology’s MLST 2.0 ^25^, ResFinder 4.0 ^26^, PlasmidFinder 2.1 ^27^, MGE (mobile genetic element) ^28^, and Restriction-ModificationFinder 1.1 ^29^. The GenBank annotation files were downloaded and parsed through SnapGene Version 7.2.1 to illustrate the genetic environment of the resistance genes.

### Phylogenomics

*A. baumannii* strains that were classified as resistant by computational means or through laboratory analyses were selected from the PATRIC database. Those from Africa were separated and downloaded and those from other continents were also grouped together: a selection of 199 strains were randomly obtained from the non-African group and the genomes from the African strains were downloaded. Finally, sections of the strain’s genome was BLASTed to identify other strains that closely aligned with it. The genomes of these closely aligned strains through BLAST analysis were also downloaded for additional phylogenetic analysis. The downloaded genomes, together with this study’s genome, were aligned using ClustalW and ≥1000 coding sequences were used to phylogenetically analyse their evolutionary relationship using the randomized axelerated maximum likelihood (RAxML) tool. Default parameters were used except that 1000 genes were set as the minimum for all genomes and a bootstrap of 1000 was used. The Newick file was annotated using FigTree v1.4.4.

### Epigenomics

The Restriction Enzyme Database (REBASE) ^29^, hosted by the Centre for Epidemiology was used to identify the restriction modification system (RMS), which includes DNA methylation, restriction endonucleases, and their motifs ^30^. The methylation modifications and motifs were also determined using PacBio’s MotifMaker software ^29^. PGAP annotations of the contigs also identified the restriction endonucleases (REs), methylases or methyltransferases (MTAses), and associated methylation genes in each contig. These annotations were visualized using SnapGene 7.2.1.

### RNA-sequencing data analysis

HTSeq-DeSeq2 was used to align, assemble, and evaluate the differential gene expression of the isolate. *A. baumannii* ATCC 1909 strain was used as the reference genome. The function of each gene was evaluated using the genome annotations of the reference strain on the PATRIC database.

### Data availability

This Whole Genome Shotgun project, epigenomic, and RNAseq data have been deposited at DDBJ/ENA/GenBank under the BioProject number PRJNA861833 and accession number JANIOU000000000. The version described in this paper is version JANIOU020000000.

## 3. Results

### Identification, Typing, and Resistance Profile

The isolate was identified by the MicroScan Walkaway identification/antibiotic susceptibility testing system (Beckman Coulter Diagnostics, United States) using Panel Combo 68 as a non-fermenting, MDR and ESβL-producing isolate. The species was confirmed by NCBI’s ANI ^24^ to be *A. baumannii*, with resistance to 22 out of the 25 antibiotics tested: amikacin, amoxicillin-clavulanate, ampicillin/sulbactam, ampicillin, aztreonam, cefepime, cefotaxime, cefoxitin, ceftazidime, cefuroxime, cephalothin, ciprofloxacin, colistin, ertapenem, fosfomycin, gentamicin, imipenem, meropenem, norfloxacin, nitrofurantoin, tobramycin, and trimethoprim-sulfamethoxazole. It was however sensitive to tigecycline and piperacillin-tazobactam, and had intermediate susceptibility to levofloxacin and nalidixic acid. The BMD and disc diffusion tests confirmed isolate R-B2.MM as resistant to both colistin (>128 µg/mL) and the carbapenems (zone diameter >19 mm).

The multiplex PCR screening of the isolate detected *mcr-*1 and a *bla*_OXA_-like gene, while the whole-genome sequencing identified 27 resistance genes (or 31 resistance genes with variants): *aadA1* (six copies: three copies on chromosomal contig 1 and three copies on chromosomal contig 2), *aac(3)-Ia, aph(3)-Ia, aph(3’’)-Ib, aph(6)-Id* (two copies on the chromosome; contig 1), *armA, arr-2, dfrA1* (four copies: two copies on chromosomal contig 1 and two on chromosomal contig 2), *dfrA15, mcr-1.1* (two copies on plasmid contig 3), *mphE, msrE, qacE, qnrS1, sitABCD, strA, strB, sul1* (three copies on chromosomal contig 1), *sul2* (three copies on chromosomal contig 1), *sul3, cmlA1* (two copies: one on contig 1 and another on plasmid contig 3), *qnrS1, tet*(B), *tet*(M), *tet*(A) (two copies on plasmid contig 3), *bla_PER-7_, bla_OXA-69_, bla_OXA-23_* (three copies on chromosomal contig 1), *bla_ADC-25_*, *bla_TEM-1B_*, and *bla*_NDM-1_. The resistance phenotype corresponded to the resistome in that the resistance genes identified corresponds to the resistance profiles observed from the MicroScan and BMD (Table S3: Dataset 1).

The strain was found to belong to multilocus sequence types (MLSTs) ST1604, ST231, or ST1: ST1604 and ST231 from the Oxford MLST scheme and ST1 according to the Pasteur MLST Scheme. This strain contained 12 virulence genes: cma, ompT, traT, iutA, iucC, iss, hlyF, iroN, sitA, cib, cvaC, and ipfA.

Among the four EPIs, EDTA (71.3-fold change) and CCCP (2.5-fold change) had a significant MIC reduction (≥ 2-fold) on colistin while the rest did not. Notably, the *P. aeruginosa* ATCC 27853 had a significant reduction in colistin MIC in the presence of EDTA (2-fold reduction) and reserpine (4.7-fold reduction). Contrarily, only EDTA caused a reduction (22.7-fold) in the MIC of ertapenem while PAβN and CCCP significantly reduced the MICs of ertapenem by 2-fold in the *E. coli* ATCC 25922.

### Genetic environment of resistance genes

Contigs 1 and 2 were chromosomal, with contig 1 containing more resistance genes than contig 2. Notably, contigs 1 and 2 also had their resistance genes being clustered together in one region flanked by integron cassettes and composite transposons, forming a genomic island. On contig 1, the resistance genes clustered between 0 – 80 kb within integrases, recombinases, insertion sequences, transposons, DNA (cytosine) methyltransferases and methylases (Fig. S1-S5). Hence, all the resistance genes within this ∼80kb genomic island were bracketed by composite transposons that also included class1 and class 2 gene cassettes. Within the genomic island on contig 1, *aadA, sat2*, and *dfrA1* were contiguous to each other and bracketed by Tns*D* and *IS*256 transposases. *Aph(6’)-I* was flanked by a class 2 integrase (*IntI2*) and IS*Vsa3* transposase, followed in close synteny by *sul2*, N-6 DNA methylase, IS*26*, and class 1 integrase (*IntI1*) in the reverse orientation to the *IntI*2. Following this *IntI1* integron cassette were batteries of *IS*s, transposases, and resistance genes such as *arr-2, cmlA5, QacE, sul1, bla_PER_, sul1, armA, msr(E), mph(E), tetR(B),* and *sul2*. Instructively, there were two copies of *sul1* and *sul2* genes within the genomic island (Fig. S2-S3).

Indeed, the whole genomic island of ∼80kb seems to be a composite transposon and an episome as a BLAST analysis showed that it aligned with several plasmids such as Ab04-mff plasmid pAB04-1 (CP012007.1) and chromosomes such as that of *A. baumannii* AR_0083 (CP027528.1). Moreover, replicative transposition was observed at 1.19 – 1.22Mb, 1.23 – 1.28Mb, and 1.32 – 1.33Mb regions on contig 1 (Fig. 1 and 2A) with *bla_OXA-23_*flanked by a type IIL restriction-modification enzyme *Mme*I and two IS*Aba1* transposases at both sides, forming a composite transposon. In Figure 1A, additional Tn*SA*, Tn*iB*, Tn*iQ* and Mu transposases were in close proximity to the composite IS*Aba1*-flanked transposases while a homocysteine S-methyltransferase was found in close proximity to the same composite transposon in Figure 1B. Thus, three copies of *bla_OXA-23_* within an IS*Aba1*-flanked composite transposon were present in the genome owing to this replicative transposition. These three copies ranged from 1.19 – 1.34 Mb (Fig. 1 & 2A), which aligned to both chromosomes (including that of *A. baumannii* AR_0083 (CP027528.1)) and to an unnamed1 plasmid (90% query cover and 97.7% nucleotide identity) from *A. baumannii* 2021CK-01407 (CP104448.1) when BLASTed.

**Figure 1.**
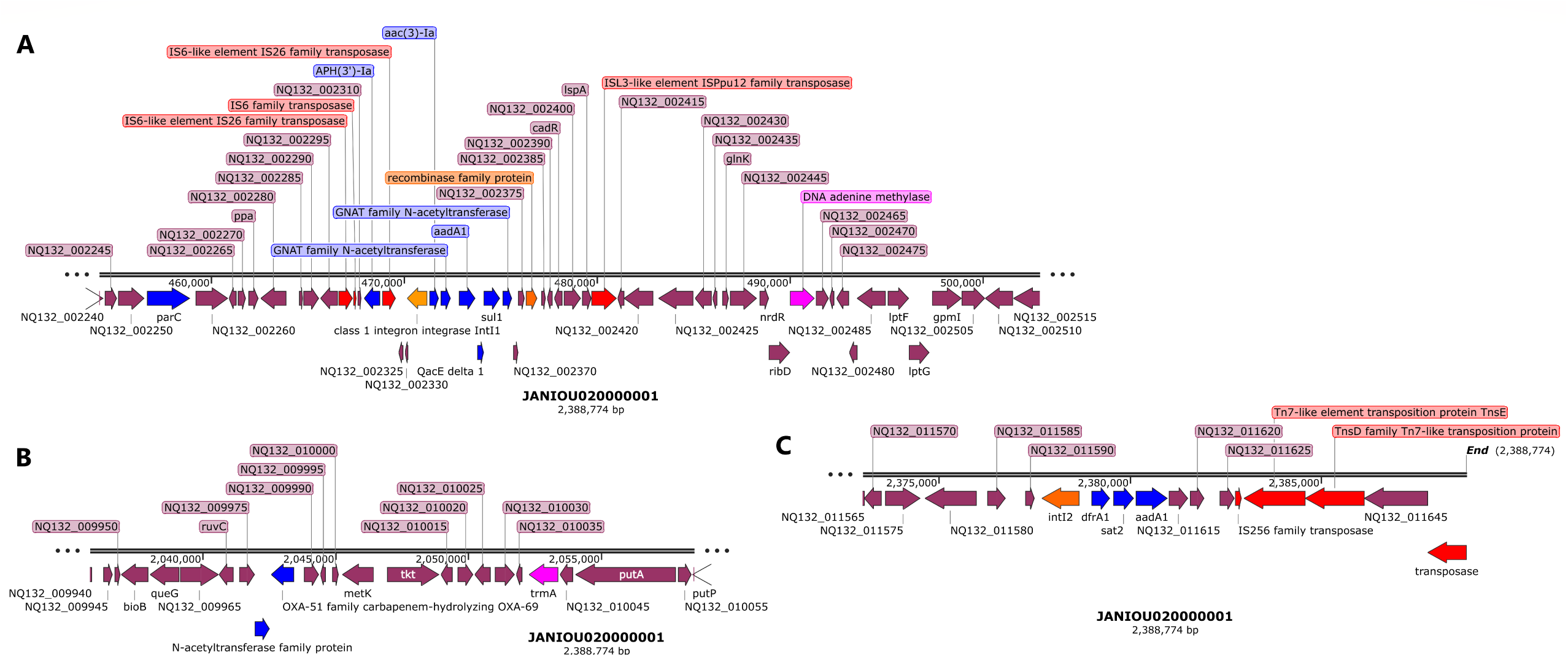
Genetic environment of blaOXA-23 carbapenemase on chromosomal contig 1 (1.19-1.22Mb, 1.23-1.28Mb). Mobile genetic elements (shown as red arrows) and methylases/restriction modification endonuclease (purple arrows) bracketing the *bla_OXA-23_* gene shows that the gene is within a composite transposon. The same chromosomal contig has two *bla_OXA-23_* genes shown in A and B, with the genetic environment in A being different from that in B. In both cases, *bla_OXA-23_* was bracketed by IS*Aba1* and transposases.

**Figure 2.**
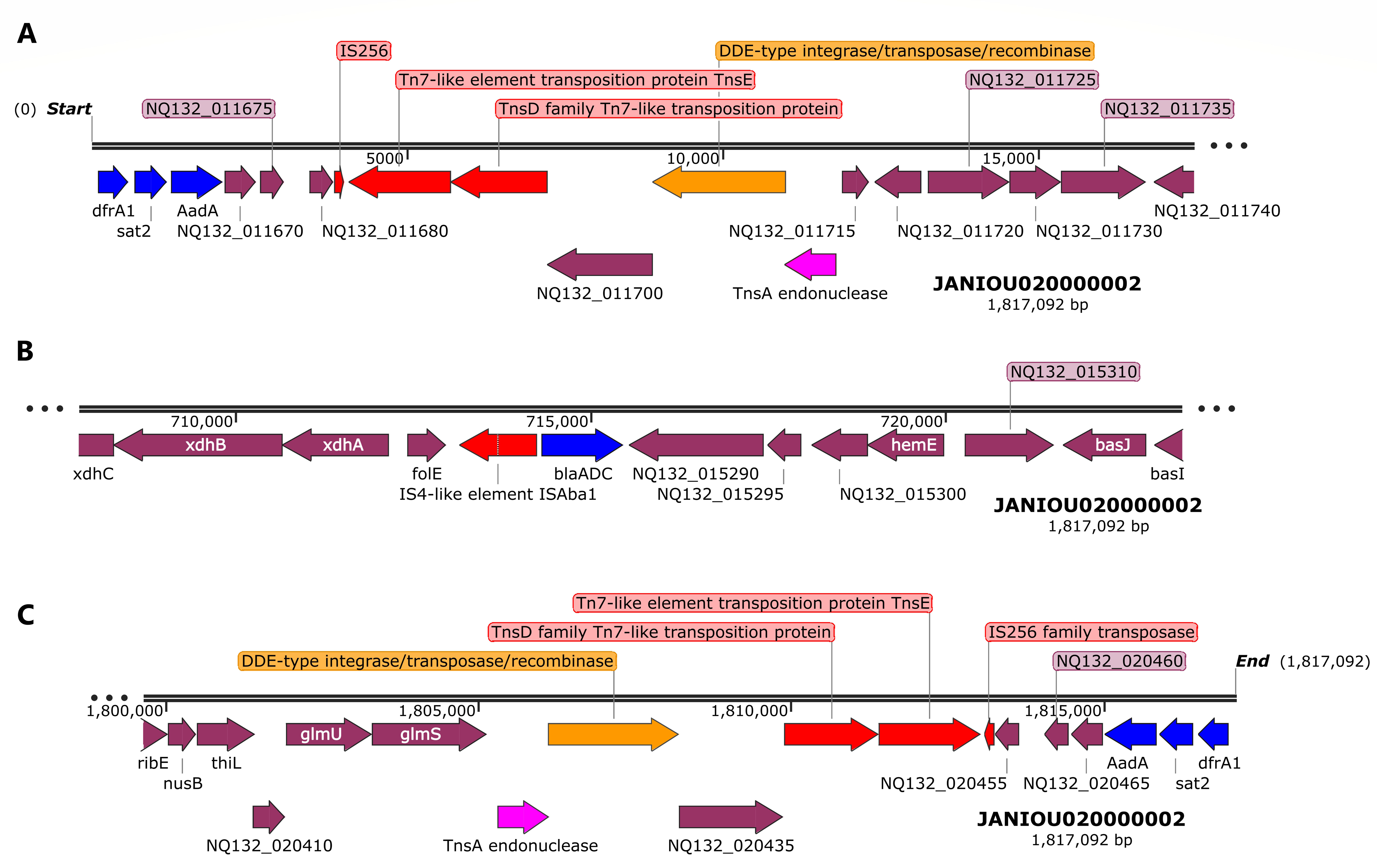
Genetic environment of sul2, aph(3”)lb, aph(6)-ld, and blaOXA-23 and blaNDM-1 carbapenemase on chromosomal contig 1 (∼1.32-1.34Mb, ∼320-360kb). *bla_OXA-23_* (shown as blue arrow) in A, is bracketed by a composite transposon consisting of IS*Aba1* and transposases. *sul2, aph(3”)lb, aph(6)-ld, and bla_NDM-1_*, in B (shown as red arrows), were sandwiched between IS*1006*, recombinase, IS*Aba1*, IS*30*, and IS*Vsa3* insertion sequences and transposases.

Another genomic island with a composite transposon flanking *aph(3”)-Ib, aph(6)-Id*, *bla_NDM-1_*:*ble*, and *sul2* was found between 328 – 353kb on contig 1, which aligned to both *A. baumannii* plasmids and chromosomes: CP027528.1, AP031576.1, CP130627.1, CP035935.1, CP090865.1. The composite transposon comprised of IS*1006*, recombinase, IS*Aba1*, IS*30*, IS*91*, and IS*Vsa3* with IS*30:bla_NDM-1_:ble* synteny being present (Fig. 2B). A third genomic island was also identified on contig 1 between 455kb and 500kb (Fig. 3A), which comprised of a class 1 integron (*IntI1*), a recombinase, DNA adenine methylase, IS*Ppu12* and IS*26*-IS*6* transposases, and *aph(3’)-Ia*, GNAT N-acetyltransferase (2 copies), *aadA1, QacEΔ1, aac(3’)-Ia, aadA1*, and *sul1* resistance genes. This genomic island was in close synteny with an upstream *par*C gene and mostly aligned with chromosomes and a single plasmid (from *A. baumannii* 2021CK-01407 (CP104448.1) when BLASTed (Fig. 3A).

**Figure 3.**
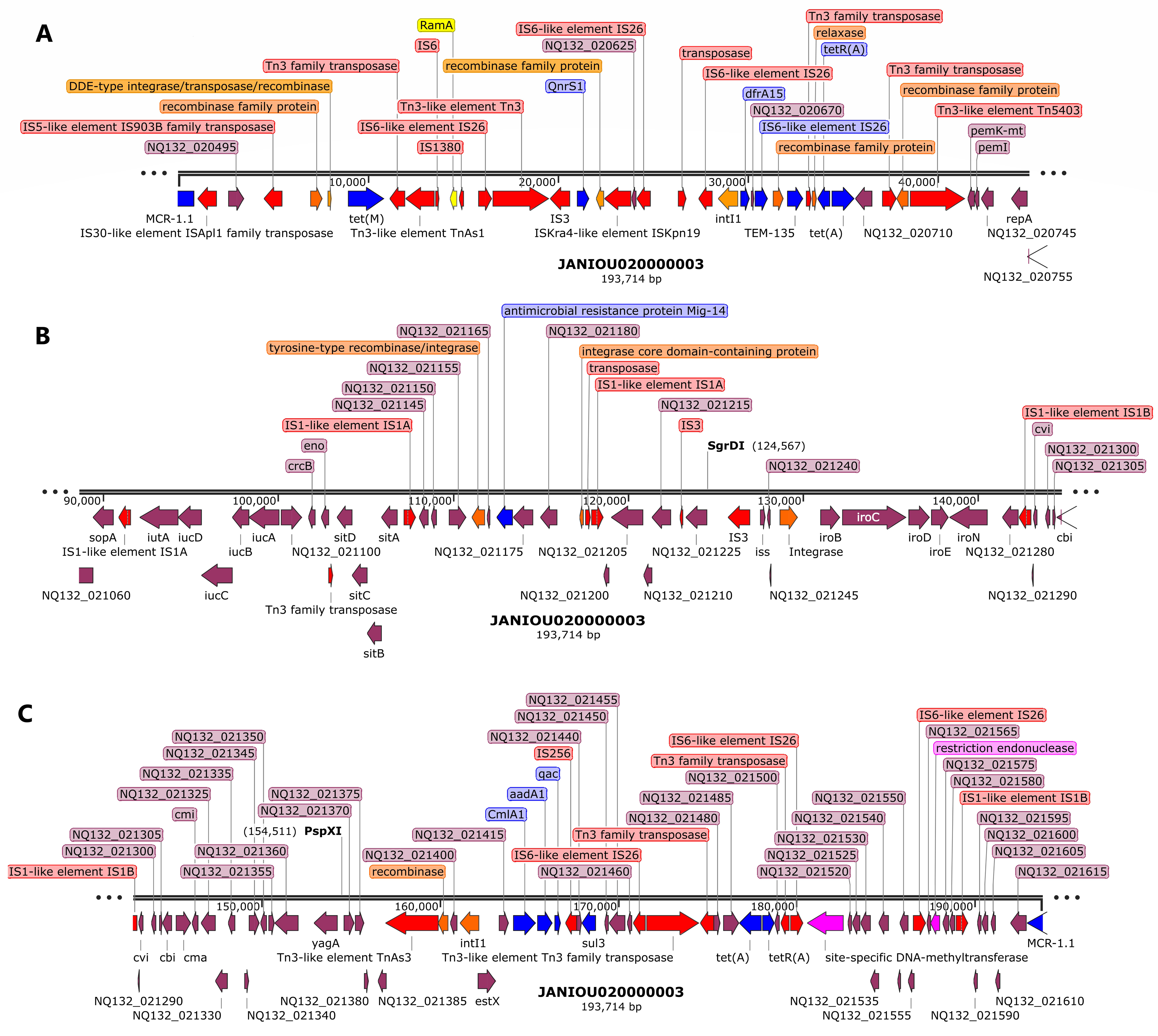
Genetic environment of aph(3’)-la, aac(3)-la, aadA1, sat2, dfrA, and Sat2 resistance genes on chromosomal contig 1 (∼450-500kb, 2.04-2.06Mb, ∼2.375-2.39Mb). IS*26*-IS*6* and IS*Ppu12* transposase bracketed *aph(3’)-la, aac(3)-la,* and *aadA* while a recombinase and class 1 integron integrase (*IntI1*) sandwiched *sul1, qacE*, GNAT family N-acetyltransferase, and *aadA1* genes. Resistance genes are shown as blue arrows and mobile genetic elements are shown as red arrows. A DNA adenine methylase (purple arrow) was also found in close proximity to the resistance genes within the same genomic island on the chromosome (A). N-acetyl transferase and *bla_OXA-51_* carbapenemase were also found in close proximity to *trmA* methylase (purple arrow) within the same region without any mobile genetic element (B). *dfrA, sat2,* and *aadA1* are also bracketed by a class 2 integron (*IntI2*), IS*256* and Tns*D/E* (Tn-*7*-like) transposases (C).

Between 135-250kb on contig 1, methyltransferases, IS*Aba1* transposase and *gyr*B were found, without any other resistance gene (Fig. S4-S5). A *bla_OXA-51_* carbapenemase gene, in close synteny with an N-acetyltransferase and a *trmA* methylase was found within 2.04 – 2.06 Mbp on contig 1 without any MGE (Fig. 3B). Towards the end of contig 1 (2.375Mbp – end), however, *dfrA1, sat2*, and *aadA1* were bracketed by an *IntI2* class 1 integron, IS*256*, Tn*7*-like (Tn*sE*-Tn*SD*) and a truncated transposase (Fig. 3C). A BLAST analysis of this region showed that it aligned with only chromosomes from both *A. baumannii* and other Enterobacterales species such as *Proteus mirabilis, Enterobacter hormaechei, Shigella sonnei, Citrobacter freundii/gillenii, Providencia rettgeri, Morganella morganii, Escherichia coli,* and *Moellerella wisconsensis*.

Figure 4 shows the resistance genes and their genetic environments on contig 2. Notably, the beginning (0 – 15kb; Fig. 2A) and end (1.8Mb – end; Fig. 2C) of this contig has the same resistance genes and MGEs but in opposite directions/orientations: *dfrA:sat2:aadA:::*IS*26:<u>Tn</u>7-*Ts*E:*Tn*7-*Ts*D::integrase/recombinase:*Tn*SA* endonuclease in the 5’-3’ direction and Tn*SA* endonuclease:integrase/recombinase::Tn*7*-Ts*D*:Tn*7*-Ts*E*:IS*26*:::*aadA:sat2:dfrA* in the 3’-5’ direction. Between these two repeated regions, ∼714 – 716kb, is the *bla_ADC_:*IS*Aba1*resistance gene and IS (Fig. 4B). A BLAST analysis of the two repeated regions shown in Figures 2A and 2C showed that both regions aligned to chromosomes of *A. baumannii* and other Enterobacterales species.

**Figure 4.**
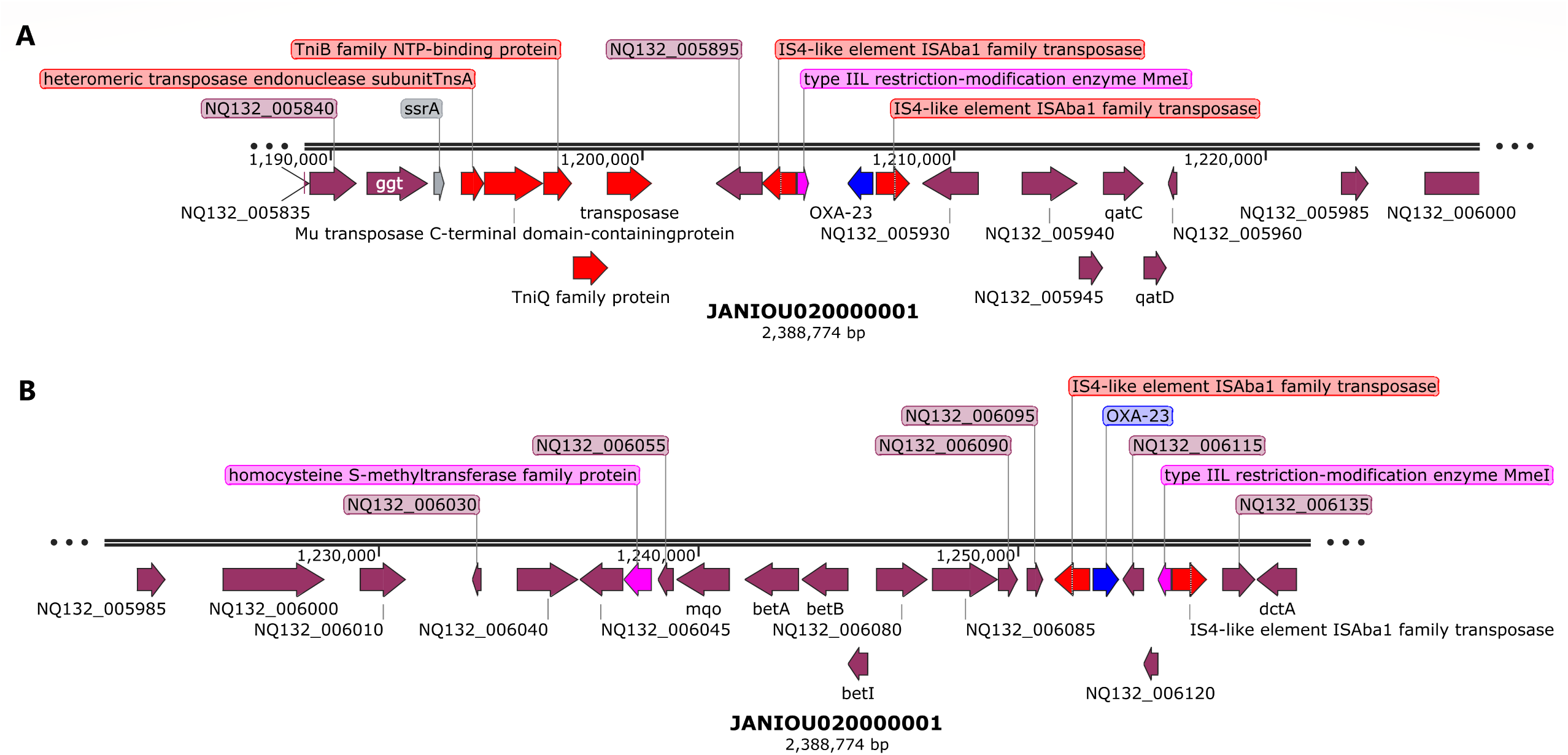
Genetic environment of aadA1, sat2, dfrA, and blaADC resistance genes on chromosomal contig 2 (0-15kb, 710-720kb, 1.8Mb-1.82Mb). IS256, Tn*sE*, and tns*D* mobile genetic elements (shown in red arrows), recombinase (orange arrow), and an endonuclease (purple arrow) bracketed *dfrA, sat2, aadA (A)* while *bla_ADC_* was in close synteny with IS*Aba1* (B). The region shown in C is a “cut and paste” transposition of the region shown in A but in the reverse orientation, suggesting that a replicative transposition event occurred within the genome.

R-B2.MM had two plasmids, contigs 3 (pR-B2.MM_C3) and 5 (pR-B2.MM_C6), identified through BLAST analyses of the contig sequences and found to be circular. While the plasmid type for pR-B2.MM_C6 was not identifiable, with only a repM replicase gene found on it, pR-B2.MM_C3 was found to contain IncFIB, IncX1 and IncFIC(FII) replicase gene sequences (Fig. 5, S6-S7). Indeed, the pR-B2.MM_C6 plasmid had 100% coverage and nucleotide identity to *A. baumannii* plasmids such as CP145435.1 and CP142898.1 (Fig. S7) while pR-B2.MM_C3 plasmid had 65-77% coverage and 99% nucleotide identity with *E. coli* and *K. pneumoniae* plasmids (Fig. S6; Dataset 1: Table S3). The BLAST analyses of pR-B2.MM_C6 shows that the plasmids it aligned 100% to were more than 17462 bp long while pR-B2.MM_C6 was 8731bp with 11 protein genes. pR-B2.MM_C3 was 193,714bp with 184 protein-coding genes and only aligned with high nucleotide homology (99%) with sections of *E. coli* and *K. pneumoniae* plasmids (Fig. 5, S6; Dataset 1: Table S3).

**Figure 5.**
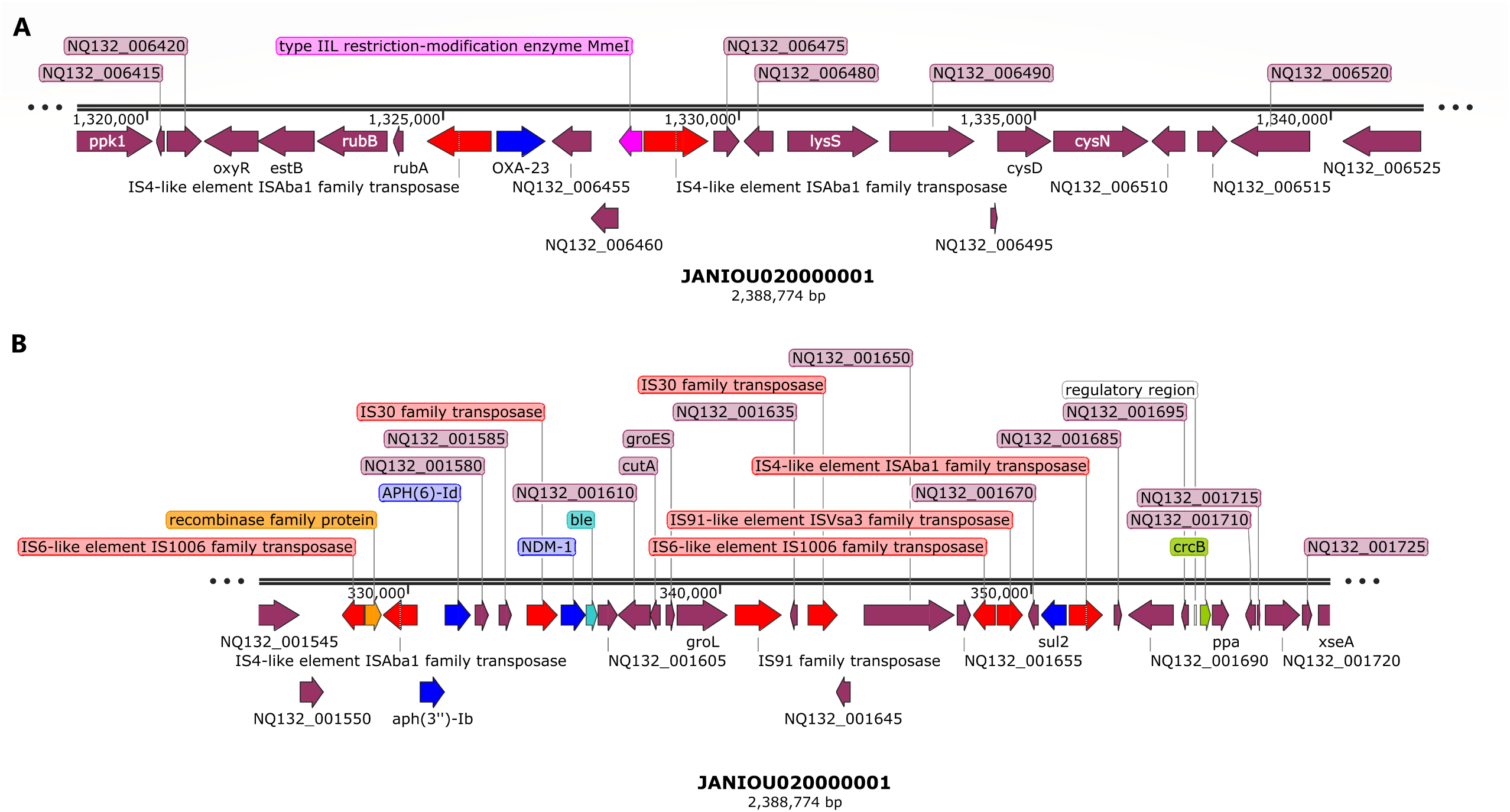
Genetic environment of mcr-1, tet(M), QnrS1, dfrA15, blaTEM-135, tet(A), cmlA1, aadA, qaC, and sul3 resistance genes on plasmid pR-B2.MM_C3 (contig 3). The resistance genes (blue arrows) were clustered together on a genomic island on pR-B2.MM_C3 at 0-42kb and 155kb-0kb and sandwiched between composite transposons (red arrows) and integrons (orange arrows). The mobile genetic environment comprised of composite transposons such as IS*903B,* IS*26,* IS*1380,* IS*1A, and* Tn*3*, alongside recombinases and class 1 *IntI1* integron. Site-specific DNA methylases were also found within this genomic island.

*AadA1, bla_TEM-1B_, mcr-1, cmlA1, qnrS1, sul3, tet(M), tet(A),* and *dfrA* were found on pR-B2.MM_C3 while pR-B2.MM_C6 had no resistance gene. The resistance genes on pR-B2.MM_C3 were localized together in a genomic island (between 0 - 42kb and 155 - 0kb) (Fig. 5 & S6) with *mcr-1* being flanked by IS*Apl1* and IS*903B* transposase. *Tet*(M) and the other resistance genes on this plasmid were also flanked by a class-1 integron and recombinase, which were also bracketed by batteries of Tn*3* composite transposons and insertion sequences (ISs) (Fig. 5A & 5C). Within this genomic island were site-specific MTAses and restriction endonucleases (REs) (Fig. 5; S6). Although mobile genetic elements (MGEs) such as IS*1A*, IS*IB*, and IS*3* were also found within the 90kb – 150kb region, only an Mig-14 resistance gene was found in this region (Fig. 5B).

### Phylogenomic analysis

The strain was not significantly related to any of the antibiotic-resistant strains from Africa used in the phylogenomic analysis (Fig. 6). It was closely related to two strains (with a bootstrap value of 5): *A. baumannii* 13367 and 13259. Expectedly, these two strains had different STs from each other and from R-B2.MM. There was also little uniformity in their resistomes, with *A. baumannii* 13367 having more similarity to R-B2.MM’s resistome (Fig. 6). Among the global *A. baumannii* strains, however, there were significant evolutionary relationship between A. baumannii strains Ab905, Ab241, Ab238, A3232, and AbCTX19 (Fig. 7) as confirmed by the high bootstrap value of 100. *A. baumannii* Ab905 and Ab241 strains isolated from blood from Israel had the same resistome, but the remaining strains had similar but not the same resistome. Notably*, bla_ADC_, bla_OXA-71/69_, bla_OXA-23_, sul1,* and *gyrA* S81L mutation was common among most of the strains. *A. baumannii* strains A3232 (Greece), AbCTX19 (France), and R-B2.MM had the same ST and belonged to the same clone.

**Figure 6.**
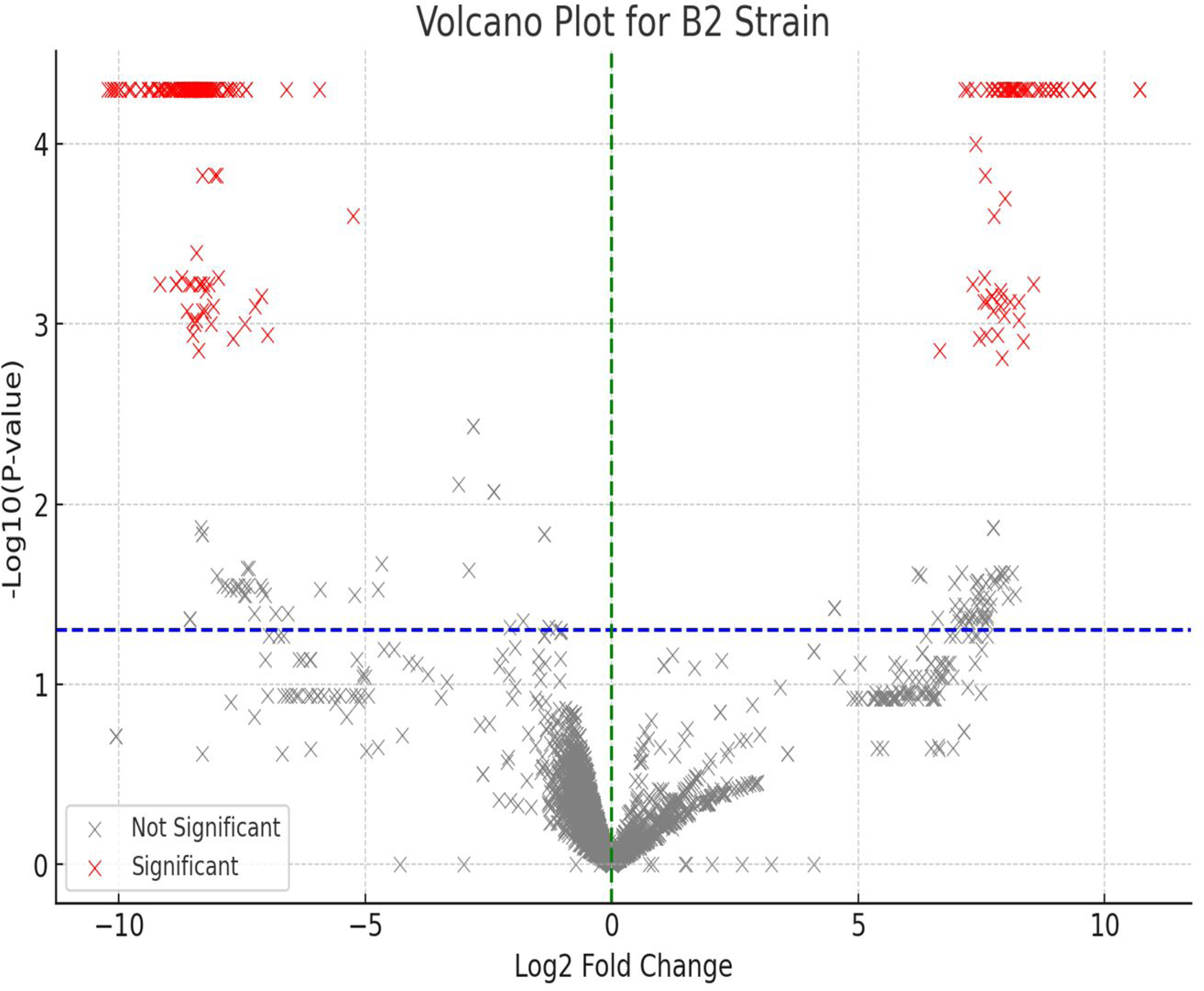
Phylogenetic analysis of antibiotic-resistant Acinetobacter baumannii strains from Africa. The R-B2.MM strain was not closely related to any resistant strain in Africa. The most closely related strains (*A. baumannii* 13259 and 13367) were not supported by the bootstrap values to be significant. The R-B2.MM strain is shown as red while all other strains are shown as black. Bootstrap values of ≥ 50 is significant. The resistomes of the three strains are also shown in a Table below the tree, with *A. baumannii* 13367 having more similarity with this study’s strain. MLST (1) is the Pasteur Institute typing scheme while MLST (2) is the PubMLST typing scheme.

**Figure 7.**
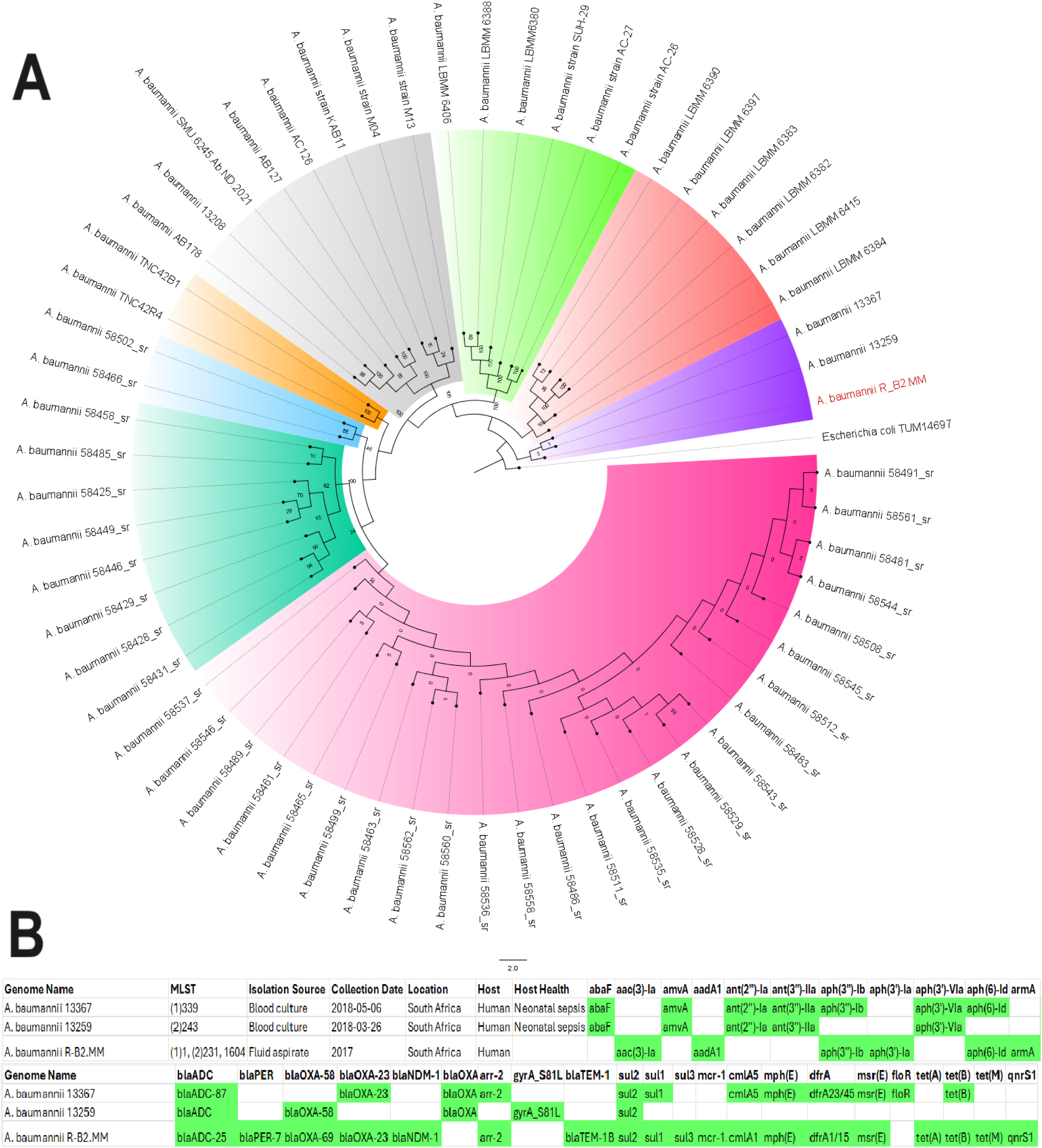
Phylogenetic analysis of antibiotic-resistant global Acinetobacter baumannii strains. *A. baumannii* Ab905 (from Tel-Aviv, Israel in 2019), Ab241 (from Tel-Aviv, Israel in 2019), Ab238 (from Tel-Aviv, Israel in 2019), AbCTX19 (from Le Kremlin Bicetre, France in 2019), and A3232 (Greece, 2022) strains formed the same clade with the R-52.MM strain. All these strains were isolated from humans and mostly from blood except strain AbCTX19 (rectal swab). Ab241 and Ab238 were of the same MLST (106 or 3) while AbCTX19 and A3232 were of MLST 231 (and 1 or 160, respectively). The significance of the clade was confirmed by the bootstrap. A3232 was most closely related to R-B2.MM evolutionarily and is thus shown as red on the tree. Bootstrap values of ≥ 50 is significant. MLST (1) is the Pasteur Institute typing scheme while MLST (2) is the PubMLST typing scheme.

*A. baumannii* strains that aligned closely with this study’s strain, with more than 80% nucleotide homology and 70% query length with sections of R-B2.MM genome, were used to conduct a phylogenomic reconstruction analysis. As shown in Figure S9 (Dataset 2), 33 strains had very close evolutionary relationship with this study’s strain, most of which was of ST231 or ST1 clone. Although these strains were isolated from different sources, different years, and different countries, they were closely related to each other. However, the resistome was not conserved across these strains.

### Epigenomics

REBASE identified types I (M.Aca7364II and M.Aba0083I) and II (M.Aba858II) methyltransferases (MTAses), with types III and IV being absent: both types were found on only contig 1 (chromosome). A specificity subunit (S.Aba0083I) was also found on contig 1 (Dataset 4). The recognition sequence of these REs were different except for only M.Aba0083I and S.Aba0083I. GATC motif was identified by PacBio with N6-methyladenosine (m6A) modifications (Dataset 4). Type IIL restriction modification enzyme MmeI, homocysteine S-methyltransferase family protein (Fig. 1-2), Tr*mA* and DNA adenine methylase (Fig. 3), Tn*sA* endonuclease (Fig. 4), and site-specific DNA-methyltransferase and RE on pR-B2.MM_C3 contig 3 (Fig. 5) were annotated throughout the genome. The annotated MTAses and REs were mostly associated with the MGEs within the composite transposons or flanked by ISs.

### Differentially Expressed Genes

The fold changes of each coding gene’s expression levels are shown in supplementary dataset 3. Out of the 4220 coding genes, 261 were significantly expressed while 3959 were not significantly expressed. As show in the volcano plot in Figure 8 and in the summarised bar chart in Figure S8, most of the significantly expressed genes were hypothetical proteins with unknown functions, followed by LysR family transcriptional regulators, phage replication proteins, class A beta-lactamases, GNAT resistance genes, outer membrane proteins, type I RMS, integrases, ABC/MFS efflux transporters, type-6 secretion systems (TSSS), OprD porins, etc. There were also MTAses, endonuclease III, MGEs, ABC/RND transporters, and prophages that were not significantly expressed (Dataset 3). Notably, the following resistance genes were significantly highly expressed: *bla_NDM_, QacE, sul2,* recombinase, aminoglycoside 3’-phosphotransferase, *aph(3’)-III/aph(3’)-IV/aph(3’)-VI/aph(3’)-VII*, aminoglycoside 3’’-phosphotransferase, *aph(3’’)-I*, macrolide 2’-phosphotransferase *mph(E)/mph(G)* family, and a class A beta-lactamase.

**Figure 8.**
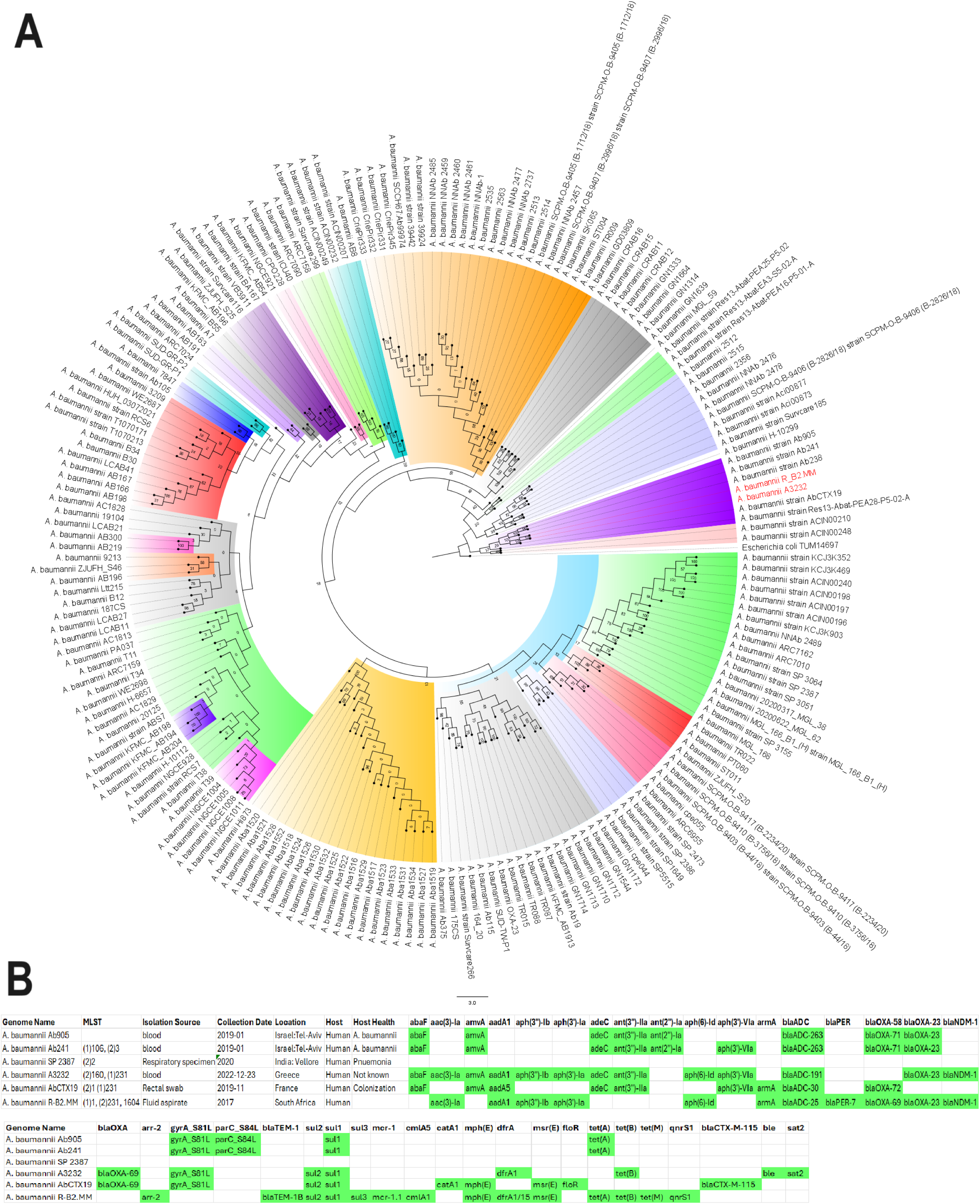
A volcano plot showing differentially expressed genes (DEGs) in Acinetobacter baumannii R-B2.MM exposed to colistin and carbapenems. The significant DEGs are shown in red while the non-significant DEGs are shown in grey. The significant DEGs include genes within the category of hypothetical protein, LysR transcriptional regulator, phage replication protein, aliphatic sulfonate monooxygenase, aldehyde dehydrogenase. Putative lipoprotein, urea carboxylase-related aminomethyltransferase, ribulose-5-phosphate 4-epimerase, and VgrG protein.

## 4. Discussion

We report on the transcriptome, resistome, mobilome, epigenome and evolutionary biology of a pandrug-resistant *A. baumannii* clinical strain that harboured NDM, OXA-23, OXA-69, ADC, MCR-1, and a plethora of resistance genes. To our knowledge, this is the first CRAB strain to co-harbour three carbapenemases i.e., NDM, OXA-23, and OXA-69, and an MCR-1 globally ^19,31^, with the *mcr*-1 being found on a plasmid. Although NDM and OXA-23 were not found on plasmids, they were found within composite transposons within sections of the chromosome that aligned with plasmids, suggesting that they might have been integrated into the chromosomes from plasmids to form episomes. This suspicion is confirmed by the replicative transposition of the composite transposon genomic islands to other loci of the genome to form multiple genomic copies of the same genes or composite transposons (Fig. 1-4). It is further corroborated by the alignment of these genomic islands to Enterobacterales species such as *E. coli, K. pneumoniae, Serratia sp., Providencia sp.,* etc. Indeed, *mcr-1* was also found within a composite transposon genomic island that can also be transposed from the plasmid to the chromosome (Fig. 5).

We are therefore of the opinion that this strain’s genome underwent multiple replicative transposition events mediated by the composite transposons, recombinases and integrons that blanketed the resistance genes, resulting in the duplication of resistance genes across the genome. The strain’s chromosome also contains several plasmid-integrated regions or resistance genomic islands, forming episomes that also aligns to other Enterobacterales species and plasmids. The evolutionary biology and phylogenetic analysis of the strain, as well as the comparative resistome analysis with other closely related strains and clones show that the resistome of our strain is quite unique (Fig. 6-7; S9). Hence, it is obvious that the genomic rearrangements observed in this genome is not wholly vertically transferred although it can vertically transfer this genome to other daughter cells as it multiplies. It can also easily transfer these resistance genomic islands horizontally to other cells through the multiple MGEs such as IS, transposons, recombinases, integrases, and plasmids found in the genome (Fig. S1-S7; Fig. 1-5).

Instructively, there was no strain from South Africa and Africa, published to date, that was closely related to this study’s strain while international strains were found to be within the same clade and of the same clone as this study’s strain (Fig. 6-7; S9). This suggests that this strain might have been imported from abroad. Notably, the other international strains that were found to be of the same clone and clade as R-B2.MM were isolated from different clinical sources such as blood, wound, urine, respiratory specimen, rectal swabs, and fluid aspirates, from different time periods spanning 1982 to 2023, and from different countries located across all continents: South and North America, Asia and the Middle East, Europe, and Australia. While this speaks to the wide geographical distribution of these strains and clade, and portend the worrying spread of MDR *A. baumannii* strains, their non-uniform resistome is also revealing. As discussed in the previous paragraph, the non-homogeneity of resistomes across the clade shows independent evolution of resistance traits as the strains spread across the globe.

The intra-genomic evolution seen in our strain, therefore, suggests that its exposure to antibiotics during treatment might have induced the replicative transposition events observed in the genome. This further supports the need to be measured in antibiotic administration to reign in the evolution and dissemination of antibiotic resistance within and across strains and species. Indeed, the transcriptomic data lends further evidence to this assertion in that we observed significant hyperexpression of genes involved in antibiotic resistance, including resistance genes, MGEs, MTAses, REs, transcription factors, membrane-associated proteins, T6SS, phage-associated genes, etc. (Dataset 3). Evidently, the exposure to antibiotics do not only cause an increase in resistance genes expression, but also in selected efflux pumps, regulatory genes, and MGEs that can both help the cell to survive by expelling xenobiotics as well as adapt its genome through accelerated transposition and transcription events to confer resistance.

This argument is supported by the presence of RMS (MTAses and REs) within the resistance genomic islands or in close synteny to the composite transposons. Given the important regulatory role of RMS in initiating or inhibiting transcription of key genes through DNA methylation, it is evident that its close association with the MGEs and its significant expression support its involvement in the observed transposition events and resistance. It has already been proven that exposure to low dose antibiotics, which lead to antibiotic-resistance mutations and adaptations, are mediated epigenetically through the RMS ^32,33^. Hence, the unique genomic arrangements and MGEs found in this isolate, which makes it different from other clones within the same clade, could most likely be mediated epigenetically. Adaptive resistance, which is mediated by epigenetic RMS factors, is transient nature and tends to disappear in the absence of the triggering factor. This further supports our assertion that this strain independently developed its MGE-mediated genomic evolution from antibiotic therapy using the epigenetic RMS pathways to regulate transcription of key genes ^32,33^.

Phenotypically, not all the EPIs resulted in a reduction in MICs of both ertapenem and colistin. Whereas EDTA and CCCP reduced colistin’s MIC, EDTA alone could reduce the MIC of ertapenem. This is expected as NDM and MCR-1 are zinc-based metallo-β-lactamases, meaning that NDM and MCR-1 cannot function enzymatically without zinc ^34^. Hence, the ability of EDTA to chelate zinc will prevent NDM and MCR-1 from being able to respectively hydrolyse their substrate antibiotics or transfer a phosphoethanolamine (PEtN) residue to lipid A in the outer membrane ^35,36^. The ability of CCCP, on the other hand, to inhibit colistin resistance is believed to be due to its ability to depolarize the plasma membrane and reduce ATP production ^35^. As colistin depolarizes the cell membrane, making it more permeable to leakages, CCCP seems to work in synergy with colistin. Hence, in the presence of CCCP, MCR-1 may be unable to counter the effect of colistin by adding PEtN to lipid A ^35^. Furthermore, the inability of the other EPIs to reduce ertapenem and colistin resistance was confirmed by the transcriptomic data in which many efflux pumps were not significantly expressed. Reserpine is also known to mainly inhibit major facilitator superfamily (MFS)-type efflux pumps in Gram-positive bacteria ^37^. Hence, its inability to affect the MICs is expected. Verapamil and PAβN are respectively known to target MATE and RND pumps in Gram-negative bacteria ^37^. However, there was no significant expression of the MFS, MATE, and RND efflux pumps from the transcriptomic data, which explains why these EPIs did not have any effect on the MICs (Dataset 3).

The repertoire of resistance genes found within the genomes correlated with the resistance phenome observed in the strain, showing that the resistance genes were being expressed to confer resistance to their respective antibiotic targets. This was also confirmed by the transcriptomic data with regards to the efflux, membrane protein and resistance genes hyperexpression (Dataset 3). Further, the association of these resistance genes on gene cassettes or within the composite transposon shows that they are moved together with the MCR-1 and carbapenemases during the transposition events or horizontal gene transfer. Hence, it is expected that this pandrug-resistant strain can share its rich resistome with other commensals or pathogens within the same niche. This will become particularly so should such a population become exposed to antibiotics, which can trigger the epigenomic and transcriptional activity towards a resistant phenotype ^32,33^. One major limitation of this study was our inability to undertake a plasmid conjugation experiment to determine the transferability of the plasmid found in this strain. Yet, the clone of R-B2.MM, ST1/ST231, is found worldwide, according to the PubMLST database, in humans with a single case, ST1/ST231, collected from the environment in Croatia. All three sequence types have been reported in Africa, specifically Ethiopia, Kenya and Ghana and, only ST1 has been previously reported in South Africa (Pretoria) in 2010 ^38^.

The immediate genetic environment of the ARGs corroborated what has already been reported globally. For instance, *mcr-1* was flanked by IS*Apl1; tet*(A) and *tet*(M) by an IS*6*/IS*26*-Tn*3; sul* and *dfrA* by a class-1 integron-IS*26; bla*_NDM_:*ble, aph(6’)-Id* and *aph(3”)- Ib* by IS*Aba1-IS*91*-ISIS*30; *bla_OXA-23_* by IS-4-like IS*Aba1; and aadA1, qacE* and *sul1* by *Inti1-*IS*6* ^31,39^. The *mcr-1* was found on an IncF-IncX hybrid plasmid while most *mcr-1* genes are found on IncH and IncC ^31,40^.

The absence of Types III and IV RMSs is confirmed by other studies ^30,31^. Most of the REs and MTAses were found on the chromosomes^30^ within the genomic islands or in very close synteny to the ARGs. However, not all the MTASes and REs were hyper-expressed significantly (Dataset 4), suggesting that only a few of the epigenomic factors were triggered by the exposure to antibiotics. There were no orphan MTAses as REs were found on both the plasmid and chromosome alongside the MTAses within the genomic islands. This contrasts with other studies that found several orphan MTAses (without corresponding REs) ^30,41^. No cytosine methyltransferase (Dcm) was found in the genome, meaning that cytosine is not methylated but adenine (Dam) is methylated at the G**<u>A</u>**TC motif, resulting in a methylated adenine at the N6 position (m6A or ^6m^A). The G**<u>A</u>**TC M6A motif is ubiquitous among prokaryotes ^30,31,41^.

Whereas some MTAses and REs were upregulated, some were not, suggesting that not all the RMSs were triggered by exposure to antibiotics. However, the functional summary of the differentially expressed genes shows that there are several unknown proteins that are marshalled in the face of antibiotics exposure to protect the bacterial cell from death. The significant differentially expressed genes (DEGs) with known functions included phage proteins and MGEs, outer membrane and efflux proteins, regulatory proteins and transcription factors, lipoproteins (useful for cell membrane structures), type-6 secretion systems, and ARGs. Instructively, eighty-five genes were upregulated while 176 were downregulated, resulting in 261 DEGs, indicating that a smaller fraction of the cellular machinery (6.2%) were marshalled to deal with the antibiotic threat. None of the RNAs in the genome was significantly expressed, which could be because colistin does not attack the RNA. Yet, it is intriguing that no RNA was hyper-expressed to produce more proteins.

## Conclusion

R-B2.MM encodes 35 resistance genes and twelve virulence genes, two plasmids (one of which is a hybrid IncX-IncF plasmid), and an MGE-rich chromosome with multiple resistance genomic islands that are episomal and aligns with plasmids and other Enterobacterales species. The strain was of closer evolutionary distance to several international strains suggesting that it was imported into South Africa. However, its resistome was unique, suggesting an independent evolution on exposure to antibiotic therapy mediated by epigenomic factors and MGE transposition events. The varied mechanisms available to this strain to overcome antibiotic resistance and spread to other areas and/or share its resistance determinants is worrying. This is ultimately a risk to public health as it was susceptible to only tigecycline. There is no better argument for antibiotic stewardship than the evidence provided herein to safeguard public health.

## Acknowledgements

This work is based on the research supported wholly/in part by the National Research Foundation of South Africa under grant number: 131013. We are grateful to Dr. Busisiwe Lebogang Skosana for her assistance with the isolates’ collection.

## Transparency declaration

None

## Author contributions

MM undertook laboratory work; NMM was a co-supervisor to the study and assisted with funding; JOS designed and supervised the study, undertook data and bioinformatic analyses and visualizations, wrote and reviewed the manuscript.

## Supplemental Tables

*Supplemental dataset 1*. Datasets comprising of Tables S1 (mcr primers), S2 (antimicrobial sensitivity testing, AST, MIC results), S3 (resistome), S4 (colistin-efflux pump inhibitors results), and S5 (carbapenem-efflux pump inhibitors results).

*Supplemental dataset 2.* Datasets comprising of metadata on demographics, clones (STs), biosamples, country of isolation, year and source of isolation, and resistance genes of strains from Africa and Globally.

*Supplemental dataset 3*. Datasets of Tables showing differentially expressed or repressed genes of Acinetobacter baumannii R-B2.MM.

*Supplemental dataset 4.* Datasets on methylation data, methylases, and DNA methylation motifs.

## Supplemental Figures

*Figure S1.* Genetic environment of resistance genes found on a genomic island on a chromosome (contig 1: 0kb to 85kb). All the resistance genes (shown in blue) were bracketed by mobile genetic elements (shown in red and orange) and clustered together on the island. The close alignment of this region with other plasmids and chromosomes show that this region was a plasmid that was integrated into the chromosome. Epigenomically, DNA methyltransferases and methylase were also found within the genomic island.

*Figure S2.* Genetic environment of resistance genes found on contig 1 (between 0kb to 40kb) shows mobile genetic elements bracketing the resistance genes. This image is an expansion of Figure S1 between the 0 - 40kb region above.

*Figure S3.* Genetic environment of resistance genes found on contig 1(between 40kb to 80kb) shows mobile genetic elements bracketing the resistance genes. This image is an expansion of Figure S1 between the 40 - 80kb region above.

*Figure S4.* Genetic environment of resistance genes found on contig 1 (∼135kb-180kb) shows mobile genetic elements bracketing the resistance genes. This image is based on the region between 130kb and180kb, showing a methyltransferase to epigenetic gene regulation. *gyr*B, which can confer resistance to fluoroquinolones when there are mutations, was found in this region without any MGE around it.

*Figure S5*. Mobile genetic elements and methyltransferases on contig 1 between 120kb to 250kb region. This region harbours methyltransferases, gyrB, and an ISAba1 mobile genetic element.

*Figure S6.* Genetic environment of resistance genes found on plasmid pR-B2.MM_C3. The plasmid contains methyltransferases, composite transposons, and integrons bracketing the resistance genes, which were clustered together within a genomic island surrounded by mobile genetic elements such as integrons, composite transposons, and insertion sequences.

*Figure S7.* Genetic map of pR-B2.MM_C6. A circularized map of pR-B2.MM_C6. pR-B2.MM_C6 has no resistance gene.

*Figure S8.* Top functional categories of significantly and differentially expressed genes (DEGs) in isolate R-B2.MM. Most of the DEGs belonged to hypothetical proteins with no known function, transcriptional regulators, and mobile genetic elements.

*Figure S9*. Phylogenetic analysis of antibiotic-resistant global Acinetobacter baumannii strains with close sequence homology to R-B2.MM. A. baumannii strains that had close sequence homology to R-B2.MM after nucleotide BLAST showed very close evolutionary association with the R-B2.MM. The strains with very close evolutionary association to the R-B2.MM strain are shown as blue text in A. The resistome is shown in B. The resistome were not conserved across the strains on the same clade. Resistance genes that were conserved across the species includes the *bla_OXA-69_* and *bla_ADC_*.

